# A novel Wnt pathway orients Par complex-dependent cell polarity

**DOI:** 10.1101/2022.06.05.494812

**Authors:** Shigeki Yoshiura, Fumio Matsuzaki

**Affiliations:** RIKEN Center for Biosystems Dynamics Research, 2-2-3 Minatojima-minamimachi, Chuo-ku, Kobe 650-0047, Japan

## Abstract

The Par-complex is a highly conserved cell polarization machinery, playing critical roles in tissue development and homeostasis. However, how extracellular signals globally orient Par-dependent cell polarity in tissues remains poorly understood. We show here that four of seven *Drosophila* Wnts redundantly serve as directional cues for the Par-dependent cell polarity of neural stem cells, neuroblasts, through a novel pathway. Wnt2/5/6/D proteins, expressed in ectodermal cells, act via frizzled receptors on underlying neuroblasts to localize armadillo/β-catenin to neuroblast cytocortex facing the Wnt signal source. The localized armadillo recruits the Par complex and Pins complex to the neuroblast apical side, through the association of arm with Par3 and Inscuteable. Wnts consequently orient the cell polarity and asymmetric divisions of neuroblasts perpendicular to the epithelium, allowing nerve tissue to grow inward along the central-peripheral axis in *Drosophila* embryos. This novel non-canonical Wnt signaling pathway may regulate Par-dependent polarity in various biological contexts.

## Introduction

Tissue development is a dynamic event involving various cellular processes, such as cell division, cell shape changes, cell rearrangements, and migration, many of which depend on cell polarization. The orientation of cell polarization is guided by the global developmental field or local cues, and is crucial for tissue development and maintenance of homeostasis^1-6^. Planar cell polarity (PCP), which determines the direction along the epithelial plane as well as that of cell migration, is typically controlled by global Wnt signaling. Another universal mechanism for cell polarization is driven by the Par complex, which plays key roles in many biological processes, including the apical-basal polarization of epithelia, asymmetric division of tissue stem cells, and zygotic cleavage. However, with the exception of a few special cases, such as chemotaxis, extracellular signaling cues that regulate Par-dependent cortical polarity remain to be elucidated^7-9^.

In the *Drosophila* embryonic central nervous system (CNS), neural stem cells, called neuroblasts (NBs), are located immediately beneath the epithelial layer and undergo stereotypical asymmetric divisions controlled by Par-dependent cell polarity. NBs divide asymmetrically and perpendicularly to the epithelial layer to produce daughter ganglion mother cells (GMCs) and descendant neurons on the opposite side of the epithelial layer; thus, the CNS tissue grows toward the center of the embryo^10-13^. *Drosophila* embryonic NBs are thus a useful model for studying mechanisms underlying the regulation of stem cell polarity and division in tissue morphogenesis.

Asymmetric divisions of the NBs are cell-autonomously driven by the apically localized Par [Bazooka (Baz, Par3), Par-6, and aPKC] and Pins [Partner of Inscuteable (Pins), Gαi and Mud] complexes. The Par complex localizes cell fate determinants, such as Prospero and Numb, which associate with their cortical adapters, Miranda (Mira) and Pon, respectively, to the basal cortex of NBs^14-24^. In contrast, the Pins complex regulates the orientation of the mitotic spindle^25-28^. These two complexes are bridged by the Inscuteable (Insc) protein that binds both Baz and Pins to co-orient the cortical polarity and mitotic spindle^21,23,26,28,29^. Cell fate determinants are subsequently segregated to the daughter GMCs during NB division.

Previous studies have suggested that the overlying epithelial cells regulate NB polarity orientation in a non-cell-autonomous manner^13,30^. In dissociated culture systems, NBs attached to epithelial cells undergo directional divisions, while those removed from epithelial cells divide in random directions. The G-protein coupled receptor (GPCR) Tre1 functions in NBs to ensure the correct orientation of the NB division axis relative to the epithelial layer; however, Tre1 is dispensable for NB polarity formation. These findings suggest that epithelial cells may provide extracellular signals to NBs to orient the NB division axis perpendicular to the epithelial layer. However, the precise nature of these signals remains unknown.

## Results

### Four Wnt proteins serve as directional cues that orient NB polarity

To identify the epithelial cell signals that control the orientation of the polarized NBs, we performed loss-of-function (LOF) and gain-of-function (GOF) screens against candidate secreted proteins and transmembrane proteins expressed in early embryos. In the LOF screen, we examined the zygotic phenotypes of the available mutants and deficiencies that disrupt candidate gene functions. In the GOF screen, candidate genes were ectopically expressed in the NBs and their progenies using *Insc-Gal4*. We measured the angle of the axis of NB polarity relative to the overlying epithelial plane using Mira as a marker of NB orientation (Fig. 1a). Mira is localized on the opposite side of the apical complex in wild-type (WT) NBs. The NB orientation measured in this manner was nearly perpendicular to the epithelial layer in WT (Fig. 1b and d). In both screens (58 lines for the LOF screen and 36 lines for the GOF screen), we found five candidate lines that showed minor defects in NB orientation (Supplementary Fig. 1a). Five of these genes (shown in red in Supplementary Fig. 1a) are involved in Wnt signaling^31,32^, which prompted us to focus on Wnt proteins as candidate regulators of NB orientation.

**Figure 1.**
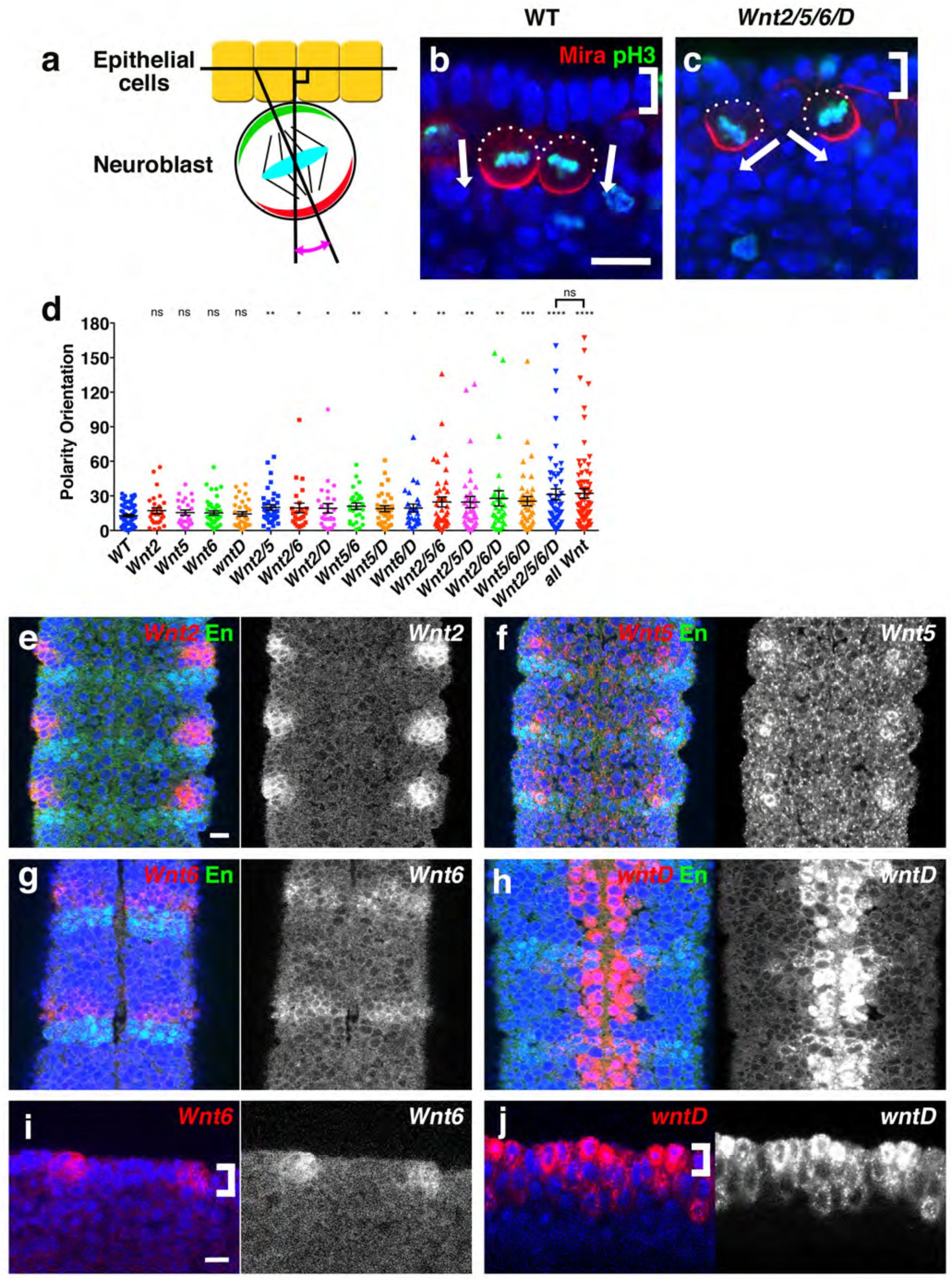
Wnt2, Wnt5, Wnt6, and wntD redundantly regulate the orientation of NB polarity. a) Method used to measure NB orientation relative to the epithelial layer. Red and green crescents indicate Mira and Par complex localization, respectively. NB orientation was defined as the arrow drawn from the midpoint of an apical component crescent to the midpoint of Mira crescent in all figures. NB orientation was measured at embryonic stage 10. (b and c) NB orientation in WT (b) and *Wnt2/5/6/D* mutant (c). Arrows: NB orientation. Brackets: epithelial layer. Scale bar: 10 μm. (d) Distribution of NB orientations in embryos for various combinations of Wnt mutants indicated below each scatter plot. each dot indicates an individual NB in the plots. Mean±SEM values are shown in the plots. ns: not significant. *: *p*<0.05. **: *p*<0.01. ***: *p*<0.001. ****: *p*<0.0001. The individual sample number n and P value (compared with WT) in the parenthesis are as follows (from left to right): *n*=70, 28 (0.081), 24 (0.27), 42 (0.26), 33 (0.49), 37 (0.0032), 24 (0.034), 27 (0.042), 27 (0.0014), 32 (0.015), 25 (0.023), 41 (0.0014), 36 (0.0030), 31 (0.0019), 42 (0.0004), 52 (<0.0001), 74 (<0.0001). The P value is 0.088 between the *Wnt2/5/6/D* mutant and the all *Wnt* mutant. (e-j) Fluorescent *in situ* hybridization showing the expression patterns of *Wnt2* (e), *Wnt5* (f), *Wnt6* (g and i), and *wntD* (h and j). (e-h): dorsal view, anterior is up. (i and j): lateral view, anterior is left. Brackets: epithelial layer. Scale bar: 10 μm.

*Drosophila* has seven *Wnt* genes (*wg, Wnt2, Wnt4, Wnt5, Wnt6, Wnt10*, and *wntD*; Supplementary Fig. 1b). Because single mutants for each *Wnt* gene showed no or very weak NB orientation phenotypes (Fig. 1d and not shown), we tested the potential redundant functions of multiple Wnt proteins in NB orientation by creating mutants that lacked various combinations of multiple *Wnt* genes. The quadruple mutant for *Wnt2, Wnt5, Wnt6* and *wntD* (designated *Wnt2/5/6/D*) showed the most defective NB orientation (Fig. 1c and d). The Par complex crescent showed highly variable orientations relative to the epithelial layer, while the alignment of the polarity axis and spindle orientation was not significantly affected (Supplementary Fig. 1c and d). Additional depletion of the *wg, Wnt4*, and *Wnt10* genes did not enhance the severity of the *Wnt2/5/6/D* mutant phenotype, indicating that these three *Wnt* genes have no role in NB orientation (Fig. 1d). We concluded that the four *Wnt* genes *Wnt2, Wnt5, Wnt6, and wntD* function redundantly in regulating the relative orientation of Par-dependent polarity in NBs.

Consistent with the idea that these four Wnt proteins are brought from the overlying epithelial cells in embryos, we found that these *Wnt* genes are mainly expressed in the epithelial cells in different but overlapping patterns at each segment (Fig. 1e-h), with weak expression in NBs, GMCs, and neurons lying beneath the *Wnt*-expressing epithelium (Fig. 1i and j and not shown). The merged expression pattern of the four genes covered nearly the entire CNS region (Supplementary Fig. 1e).

The *wg* and *Wnt4* genes also showed segmental expression patterns, while *Wnt10* showed mesodermal expression (Supplementary Fig. 1f-h, and see below). The failure of these Wnt proteins to affect NB orientation is most likely due to their secretion from the apical side of the epithelial cells, opposite to the NB side (as during segmentation), while the effective 4 Wnt proteins are perhaps secreted basally to the NB side. These results collectively indicate that Wnt signals from epithelial cells act instructively to orient NB polarity.

### Wnt signals is instructive in cell polarity orientation

The Wnt signals can be either instructive or permissive. To distinguish between these roles, we examined whether the Wnt signals from the adjacent cells are sufficient to orient NB polarity. For this purpose, we used *wg-Gal4* to drive *Wnt* gene expression in the *Wnt2/5/6/D* mutant background (Fig. 2a and Supplementary Fig. 1f). This *Gal4* line drives target gene expression in the *wg*-positive posterior line of cells at each segment, which includes both epithelial cells and delaminated NBs at this position along the anterior-posterior axis. We then measured the angle of the polarity axis of the NBs relative to the epithelial plane neighboring the *wg*-stripe cells (Fig. 2b). In the WT embryos with no transgene expression, the mean orientation of NBs adjacent of the wg-stripe was nearly perpendicular to the epithelium, with some fluctuations between –30° and 30° (Fig. 2c). This was also the case in the *Wnt2/5/6/D* mutant embryos with no transgene expression, while NBs showed more variable orientation than the WT NBs These data indicate again that wg from *wg*-expressing cells failed to alter NB orientation (Fig. 2d). In contrast, when *Wnt2* was expressed in *wg*-positive cells in the *Wnt2/5/6/D* mutant background, the average orientation of the apical polarity of the NBs was clearly tilted toward the Wnt2 source (Fig. 2e, and see Discussion). Similar results were observed when *wntD* was expressed in the wg-positive cells (Supplementary Fig. 2a). These results thus demonstrate that Wnt signals from neighboring cells are sufficient to orient NB polarity (Supplementary Fig. 2b). On the other hand, when one of the four *Wnt* genes was expressed in all NBs and NB progeny (by using *Insc-Gal4*) in the *Wnt2/5/6/D* mutant, in which condition Wnt proteins apparently surrounded individual NBs, we observed variable NB orientations, suggesting that Wnt signals are not permissive (Fig. 2f and Supplementary Fig. 2c). These results collectively indicate that Wnt signals from epithelial cells act instructively to orient NB polarity.

**Figure 2.**
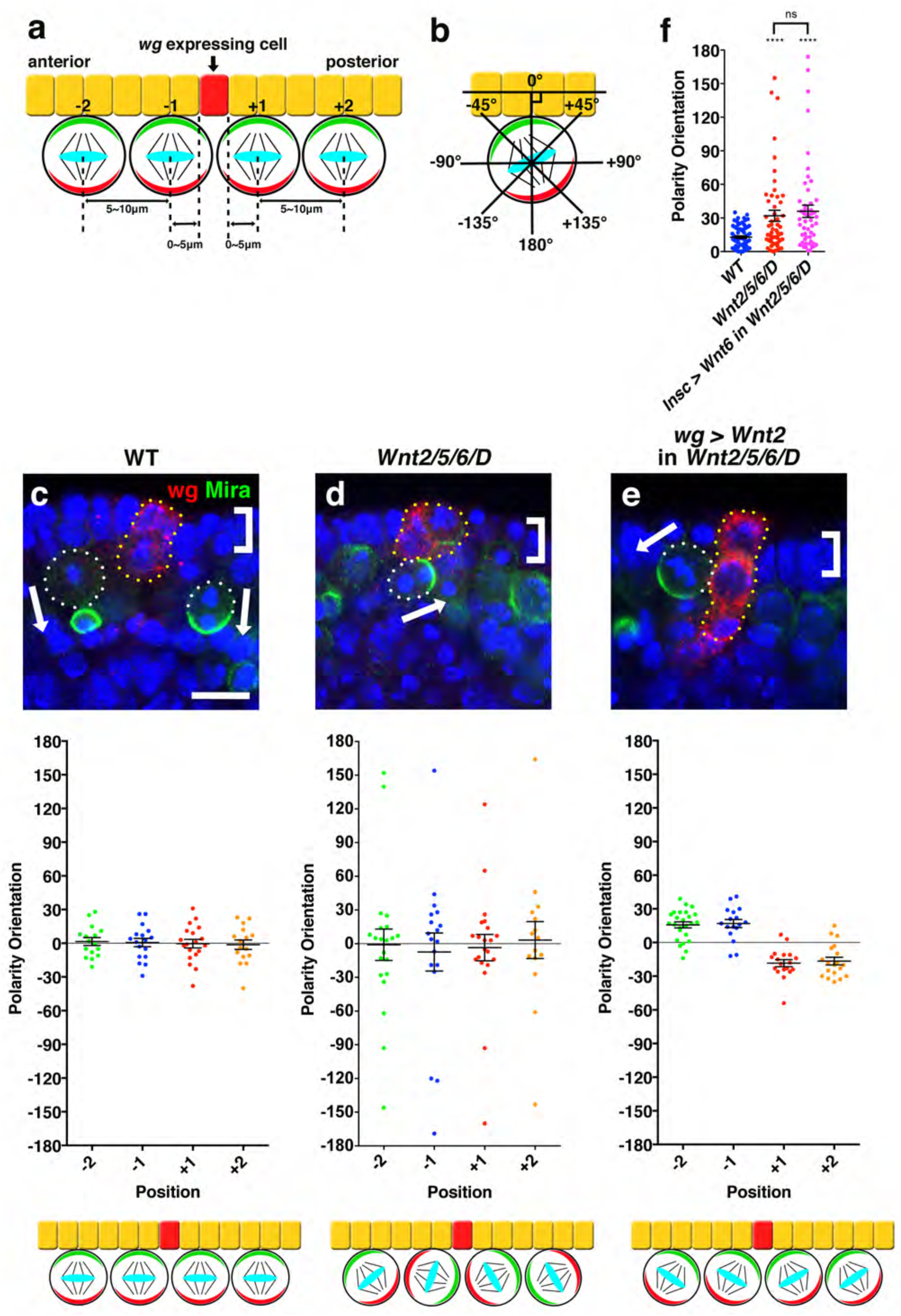
Wnt proteins from epithelial cells act as directional cues to orient NB polarity. (a and b) Method used to measure the NB orientation angle. NB positions were defined as follows: NB positions in the range of 0–5 μm anterior or posterior from the *wg* expression domain as position –1 or +1, 5–10 μm anterior or posterior as –2 or +2, respectively (a). Most NBs at position +1 and –1 are adjacent to *wg*-expressing cells, while those at position +2 and –2 are next to adjacent NBs. The angle of the NB orientation relative to the epithelial plane was measured as 0° to 180° in the clockwise direction and –0° to –180° in the anti-clockwise direction (b). (c-e) NB orientation in WT (c), *Wnt2/5/6/D* mutant (d) and *Wnt2/5/6/D* mutant expressing *Wnt2* from *wg-Gal4* (e). Top panels show a lateral view around wg-expressing cells in an embryo. Arrows: NB orientation. Brackets: epithelial layer. *wg*-expressing cells are indicated by yellow dotted lines. Scale bar: 10 μm. Middle scatter plots show the distribution of NB orientations. Mean angles are as follows: (c) position–2: 1.5°, position–1: 0.6°, position+1: –0.4°, position+2: –1.3°; (d) position–2: –1.0°, position–1: –7.4°, position+1: –3.5°, position+2: 3.2°; (e) position–2: 16°, position–1: 17°, position+1: –18°, position+2: –17°. Mean±SEM values are shown in the plots. The sample number *n*=16, 17, 19, 16, 21, 18, 21, 15, 25, 17, 18, 19 (from left to right). The bottom panels schematically show the result in each genotype. (f) Relative NB orientations in WT, *Wnt2/5/6/D* mutant embryos, and in the *Wnt2/5/6/D* mutant embryos expressing Wnt6 driven by the insc promoter that allows Wnt6 to be expressed in NBs and their progeny. Mean±SEM values are shown in the plots. ns: not significant. ****: *p*<0.0001. The sample number *n*=70, 52, 52 (from left to right). The P value is 0.58 between *Wnt2/5/6/D* mutant and *Insc>Wnt6* in the *Wnt2/5/6/D* mutant.

### fz and arm, but not pan, act as downstream effectors of Wnt2, Wnt5, Wnt6, and wntD

We searched for the downstream effectors of *Wnt2, Wnt5, Wnt6*, and *wntD. Drosophila* has four frizzled Wnt receptors (fz, fz2 to 4) (Fig. 3a)^33^. Depletion of all fz functions by expressing dominant-negative forms of fz and fz2 in *fz3/fz4* double mutant NBs compromised NB orientation similar to the *Wnt2/5/6/D* mutant (Fig. 3a). Restoring the function of one of the four fz proteins fully rescued this phenotype (Fig. 3a), indicating that all fz receptors redundantly act to regulate NB orientation in a cell-autonomous manner.

**Figure 3.**
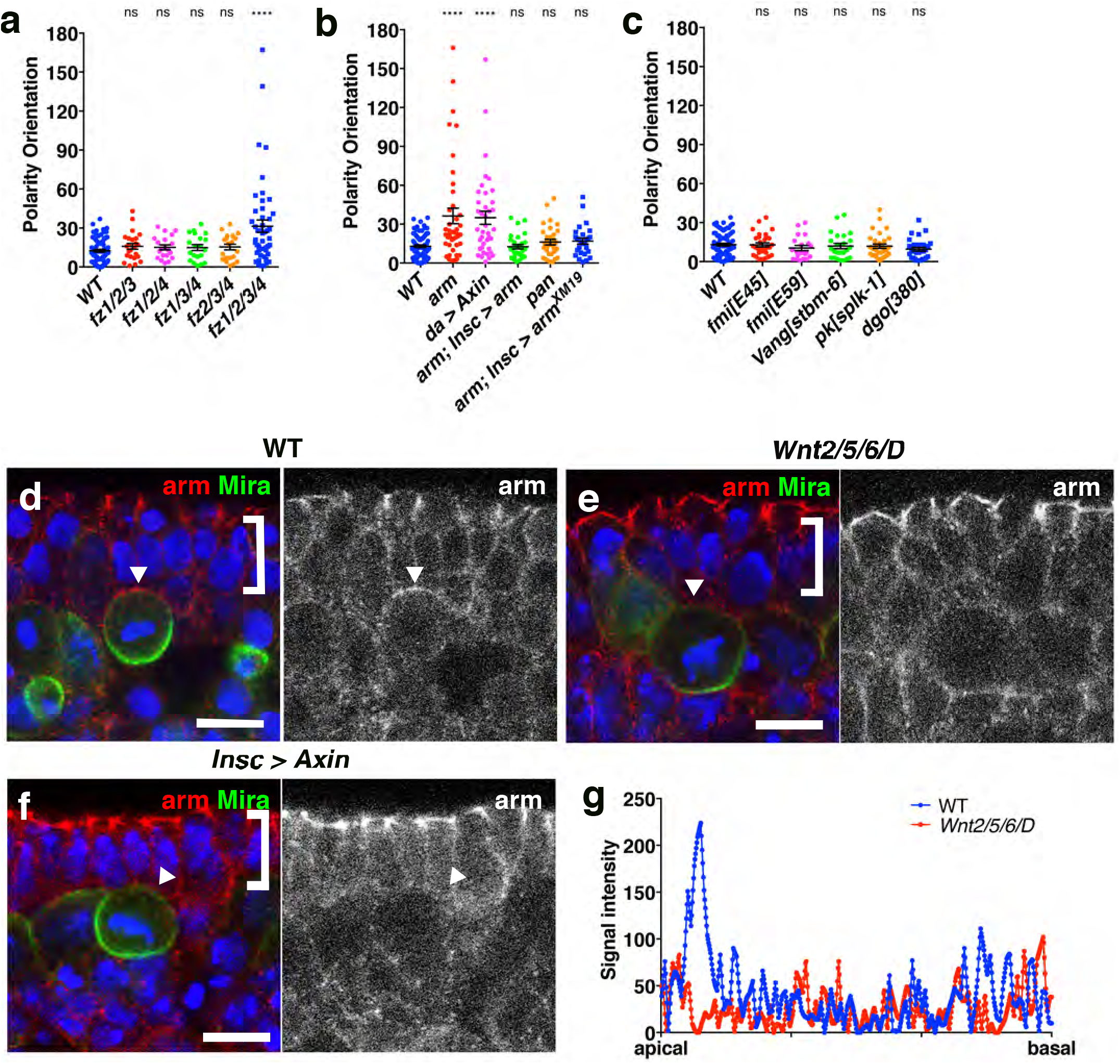
NB orientation is regulated by a novel non-canonical Wnt/fz/arm pathway that localizes arm to the apical cortex. (a-c) Statistics of the NB polarity orientation in WT and various genotypes indicated below each column (see text for details). In (b), Axin was expressed in all cells due to the use of the *daughterless* promotor. Mean±SEM values are shown in the plots. ns: not significant. ****: *p*<0.0001. The individual sample number n and P value (compared with WT) in the parenthesis are as follows (from left to right): (a) 79, 24 (0.14), 20 (0.28), 20 (0.32), 21 (0.23), 49 (>0.0001); (b) 79, 42 (>0.0001), 39 (>0.0001), 29 (0.88), 32 (0.13), 27 (0.075); (c) 78, 31 (0.93), 20 (0.26), 24 (0.63), 31 (0.54), 29 (0.076). (d-f) Localization of arm in WT embryo (d) and *Wnt2/5/6/D* mutant (e) embryos, and an embryo expressing *Axin* (driven by insc promoter) in NBs and their progeny (f). arm localization is indicated with arrowheads. Brackets: epithelial layer. Scale bar: 10 μm. (g) Line scans of the signal intensities of arm staining in WT (d) and *Wnt2/5/6/D* mutant (e) NBs along the NB orientation axis of the NBs indicated by arrowheads.

The canonical Wnt signaling pathway, the planar cell polarity (PCP) pathway, and the Ca^++^ pathway are downstream pathways of Wnt/fz signaling^34^. We analyzed the mutant phenotypes of the components of these pathways. Mutants of *armadillo* (*arm)/β-catenin*, the direct downstream target of fz, showed defective NB orientation (Fig. 3b). Furthermore, ubiquitous expression of Axin, which degrades arm even in the presence of Wnt^35^, recapitulated the *arm* mutant phenotype (Fig. 3b). In contrast, the expression of *arm* in the NBs rescued the *arm* mutant phenotype (Fig. 3b). These results indicate that arm is cell-autonomously required for correct NB orientation.

Notably, *pangolin* (*pan*)*/dTCF* mutants showed no NB orientation defects (Fig. 3b). Furthermore, the expression of *arm*^*XM19*^, which cannot interact with pan^36^, rescued the *arm* mutant phenotype (Fig. 3b). These results indicate that the Wnt/fz/arm pathway regulates NB orientation in a transcription-independent, non-canonical manner. In addition, mutants of the core components of the PCP pathway^37^, including *flamingo* (*fmi*)/*starry night, Van Gogh* (*Vang*), *prickle* (*pk*), and *diego* (*dgo*), showed no NB orientation defects (Fig. 3c), indicating that the PCP pathway is not involved in NB orientation. Taken together, our results strongly suggest that a novel non-canonical Wnt pathway regulates Par polarity-dependent NB orientation.

### Arm is recruited to the apical cortex by Wnt signals to interact with Baz and Insc

Because we found that NB polarity orientation was regulated by a novel non-canonical Wnt pathway, we investigated the relationship between arm and the apical polarity complex. We first examined the localization of the arm protein in the NBs and found that arm was localized at the apical cortex of mitotic wildtype NBs (Fig. 3d). This apical localization of arm was observed at least from prometaphase to metaphase in the WT NBs (Fig. 3d-g). This arm crescent was lost in *Wnt2/5/6/D* mutant NBs (Fig. 3e and g). Moreover, the overexpression of Axin in the NBs abolished the apical localization of arm (Fig. 3f), indicating that its apical localization is Wnt-dependent.

These results prompted us to examine whether arm interacts with any polarity proteins at the apical cortex of NB. We found that arm interacted with Insc *in vitro* and *in vivo* in addition to Baz^38^ (Fig. 4a-d). Because Baz is known to directly bind Insc^21,23^, we investigated how the binding of arm with these proteins affects their association. We found that arm facilitated the interaction between Baz and Insc (Fig. 4e and f). Taken together, these results suggest that Wnt from the epithelial cells enriches non-degraded arm at the apical cortex of NBs that faces overlying epithelial cells. These apically concentrated arm proteins subsequently facilitate the recruitment of Baz and Insc to the apical cortex and stabilize their interaction. We propose that this novel Wnt/fz/arm pathway consequently determines the relative orientation of the Par complex and activates it fully in NBs.

**Figure 4.**
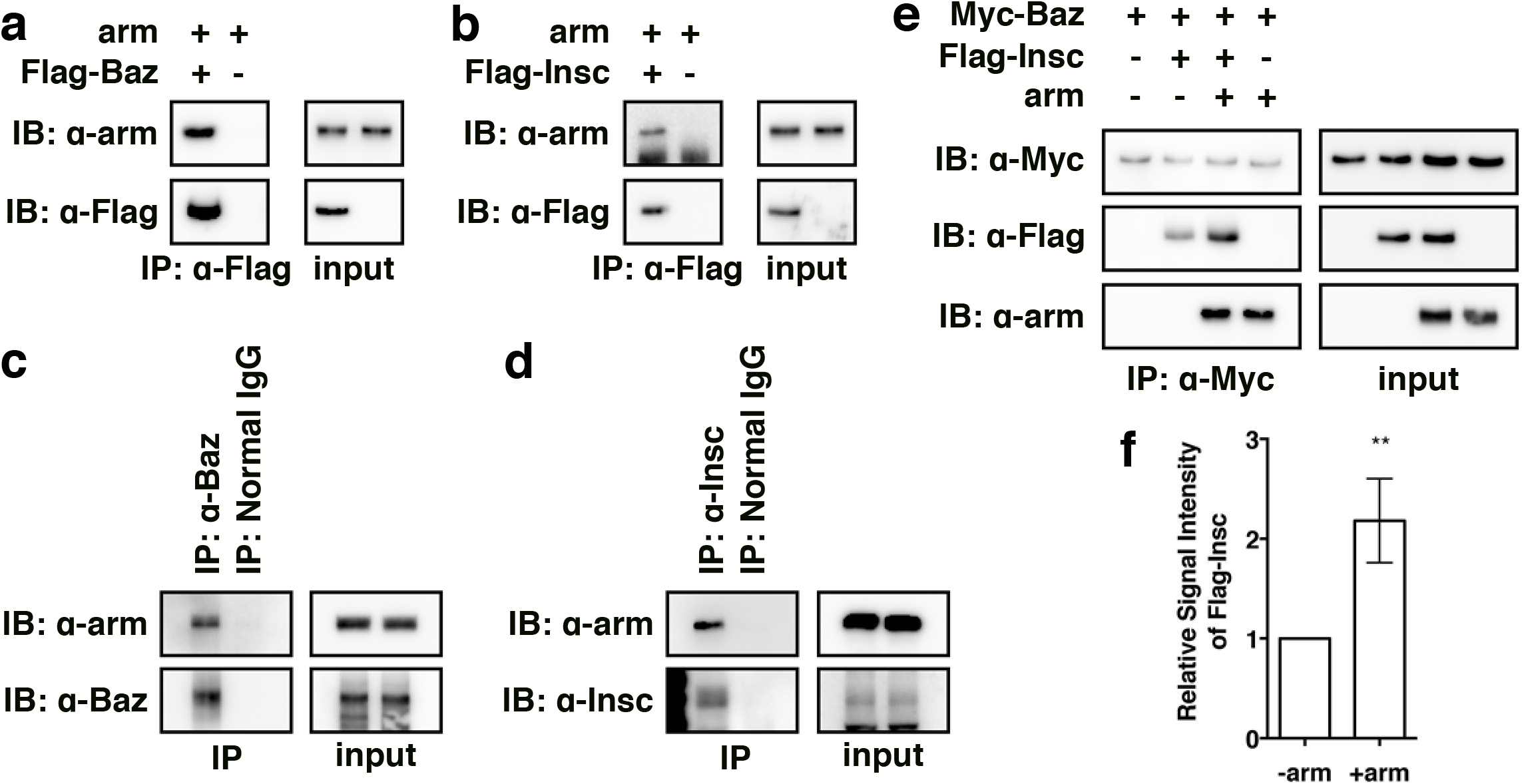
arm interacts with Baz and Insc. (a and b) Interaction between arm and Baz (a) or Insc (b) in *Drosophila* S2 cells. arm and Flag-Baz or Flag-Insc are expressed in S2 cells. The cell lysate was immunoprecipitated with an anti-Flag antibody before immunoblotting with an anti-arm or anti-Flag antibody. (c and d) Interaction of arm with Baz (c) or Insc (d) *in vivo*. Lysate of WT stage 10 embryos was immunoprecipitated with anti-Baz antibody, anti-Insc antibody, or normal rabbit IgG (normal IgG) before immunoblotting with anti-arm, anti-Baz or anti-Insc antibody. (e and f) Interaction between Baz and Insc in the presence or absence of arm in S2 cells. The cell lysate was immunoprecipitated with an anti-Myc antibody before immunoblotting with anti-Myc, anti-Flag, or anti-arm antibody. Relative signal intensities of immunoprecipitated Flag-Insc are shown in (f) (*n*=3). Mean±SD values are shown in the histogram. **: *p*=0.086.

### Wnt signaling facilitates the formation of an active Par complex

To further investigate the mechanistic basis of Wnt/fz/arm-dependent NB orientation, we examined the localization of polarity proteins in the NBs. In WT NBs, the polarity proteins localized to the apical cortex (Fig. 5a, c, e, g and i and not shown). In the *Wnt2/5/6/D* NBs, the polarity proteins, except Insc, formed a clear crescent regardless of its position (Fig. 5b, d, f, and h and not shown), indicating that the Par polarity complex itself formed almost normally. In contrast, Insc localized only faintly along the cell cortex in *Wnt2/5/6/D* mutant NBs throughout the mitotic phase (Fig. 5j). The elimination of *insc* function mislocalizes cell fate determinants, in contrast to the relatively normal localization of apical polarity proteins^21,23,29^. Therefore, it has been suggested that Insc plays a role in localizing cell fate determinants, in addition to bridging the Par and Pins complexes. We thus suspected that the faint cortical localization of *insc* in the *Wnt2/5/6/D* mutant might also result in some abnormality in the localization of the determinants in the *Wnt2/5/6/D* mutant. Indeed, we found that the formation of the Mira crescent was delayed in the *Wnt2/5/6/D* mutant. In the WT NBs, Mira started to localize basally from the end of interphase to complete the localization in prometaphase (100%, n=50, Supplementary Fig. 3a-c); the *Wnt2/5/6/D* mutant showed a diffuse Mira crescent in 76.5% of the prometaphase NBs (Fig. 5b, n=34) and in 36.7% of the metaphase NBs (Fig. 3f, h and j, n=99). In the *Wnt2/5/6/D* mutant NBs, Mira localization eventually completed before the onset of anaphase (100%, n=50). Given that the localization of cell fate determinants depends on Par complex activity, the delayed Mira localization in *insc* and *Wnt2/5/6/D* mutants is most likely caused by a delayed formation of a fully functional Par complex due to aberrant Insc distribution in both *insc* and *Wnt2/5/6/D* mutants. Thus, we suggest that Wnts ensure not only the proper orientation of NB polarity but also the efficient formation of an active Par complex.

**Figure 5.**
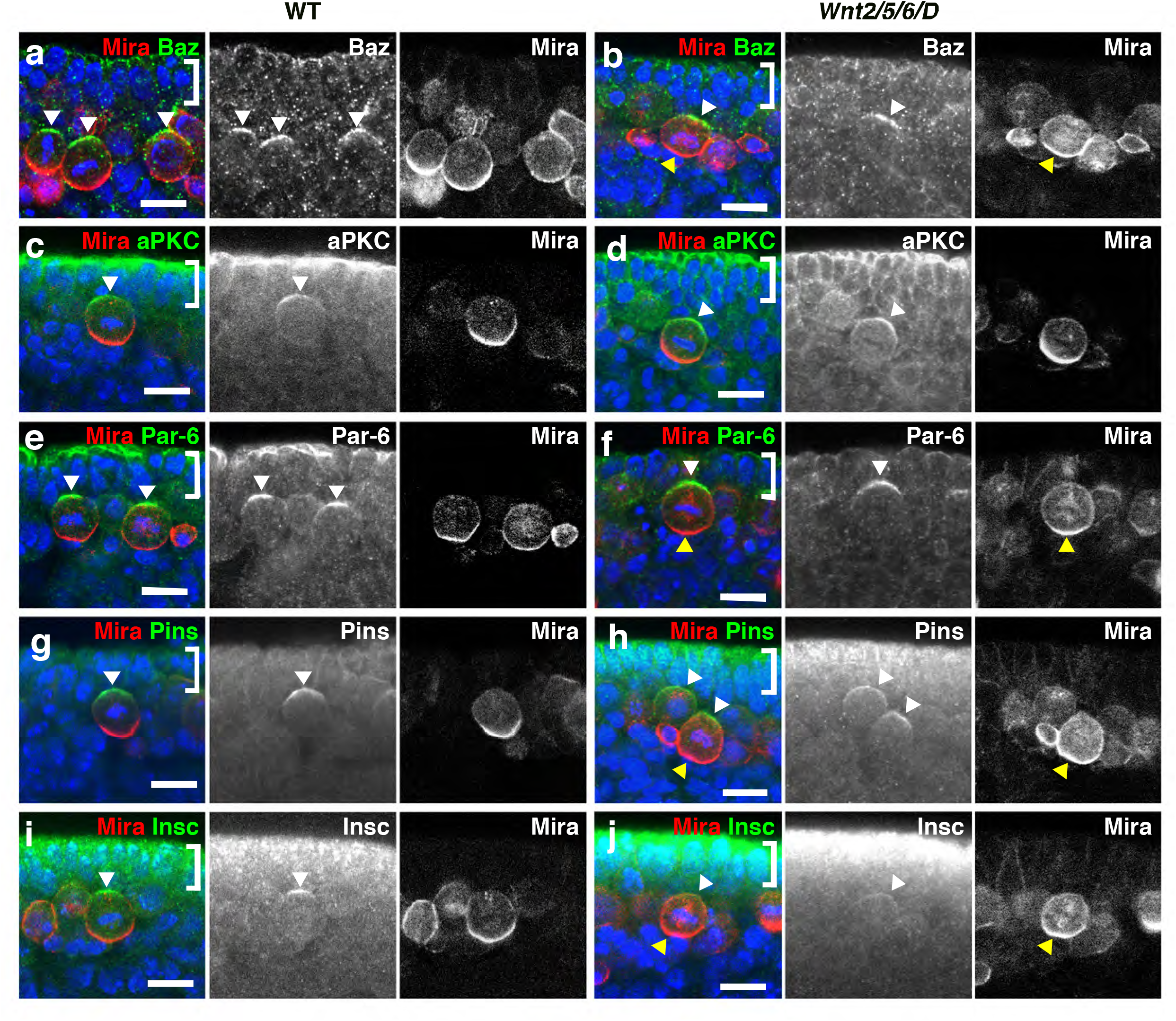
Localization of the components of the apical polarity complex in WT and *Wnt2/5/6/D* mutant. Baz (a and b), aPKC (c and d), Par-6 (e and f), Pins (g and h), Insc (i and j), and Mira (a-lj) are shown in WT (a, c, e, g and i) and *Wnt2/5/6/D* mutant (b, d, f, h and j). White arrowhead indicates polarity protein localization. Diffuse Mira localization in late prometaphase (b) or metaphase (f, h and j) NBs is indicated by yellow arrowheads. For acquiring the images of the *Wnt2/5/6/D* mutants, NBs with relatively normal orientation were selected to avoid the positional effects on the signal intensity because the background signals were higher at the apical side of the embryos. Brackets: epithelial layer. Scale bar: 10 μm.

### Wnt acts independently of Tre1 to orient NB polarity

We have previously shown that the GPCR Tre1 also orients NB polarity by enhancing Pins-Gαo interaction through Tre1 GEF function ^13^. Elimination of either of the Tre1 or Wnt pathways compromises the normal orientation of NBs. Thus, both pathways are essential for correct NB orientation. To examine the functional relationship between Wnt and Tre1, we analyzed the phenotype of a double mutant lacking these signals. The *arm tre1* double mutant displayed NB orientation defects that were more severe than those of the single gene mutants (Fig. 6a and b). Similar enhanced phenotypes were also observed when we depleted the function of all four fz receptors or expressed Axin ubiquitously in the *tre1* mutant background (Fig. 6b). Thus, the Tre1 and Wnt pathways do not function redundantly in NB orientation.

**Figure 6.**
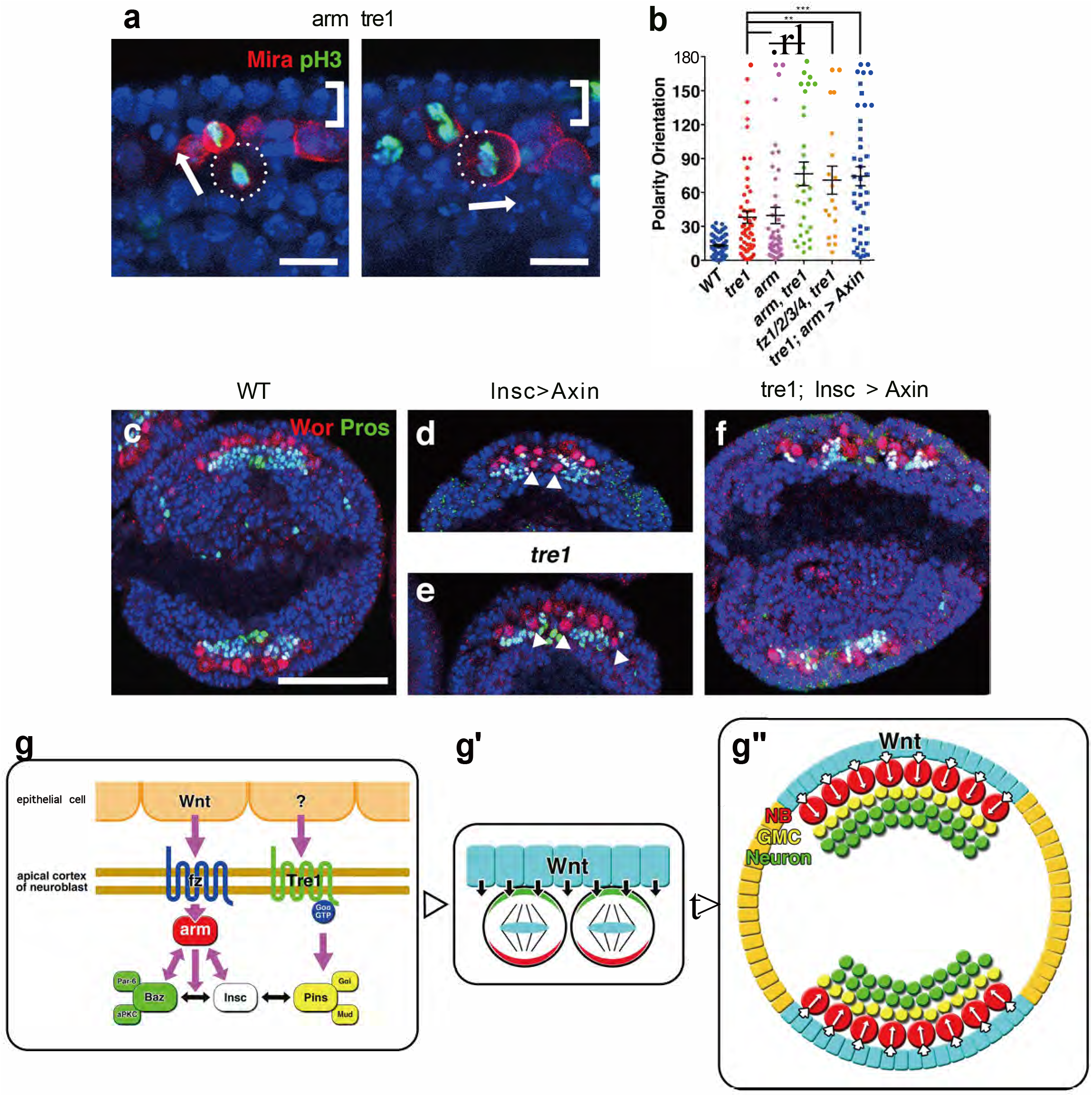
Wnt/fz/arm signal regulates NB orientation and directional growth of CNS tissue independently of Tre1 signaling. (a) NB orientation in an *arm tre1* double mutant. Arrows: NB polarity orientation. Brackets: epithelial layer. Scale bar: 10 μm. (b) Comparison of NB orientations in the absence of both *Tre1* and *Wnt* functions, using various genotypes missing *Wnt* function. Mean±SEM values are shown in the plots. The sample number *n*=84, 57, 45, 30, 19, 44 (from left to right). **: *p*<0.01. ***: *p*<0.001. The individual P value is 0.0004 for *Tre1* vs. *arm, tre1*; 0.0034 for *arm* vs. *arm, tre1*; 0.0062 for *tre1* vs. *fz1/2/3/4*; 0.0002 for *tre1* vs. *tre1, arm>Axin*. (c-f) Sections of *Drosophila* embryos showing the directional growth of CNS tissue in WT (c), embryo expressing *Axin* from *Insc-Gal4* (d), *tre1* mutant (e), and *tre1* mutant expressing *Axin* from *Insc-Gal4* (f). NBs expressed Worniu (Wor) but not nuclear Pros. GMCs expressed Wor and nuclear Pros. Neurons expressed nuclear Pros but not Wor. Arrowheads: NBs at abnormal positions. Scale bar: 50 μm. (g-g’’) Working model for Wnt/Tre1-dependent NB orientation and directional CNS tissue growth.

Next, we assessed whether the simultaneous deletion of *Wnt* and *Tre1* resulted in complete randomization of the NB polarity orientation. As shown in Fig. 6a and b, the distribution of the division angles in the absence of both Wnt and Tre1 (*tre1*; *Insc>Axin*) correlated strongly with a theoretical random distribution of the division angle (*p*=0.35, Kolmogorov-Smirnov test). Therefore, we conclude that the simultaneous loss of the two signals completely randomized the NB orientation. Together, these results suggest that Wnt and Tre1 independently act through the Par complex and Pins complex, respectively, to orient NB polarity (Fig. 6g, and see Discussion).

### Wnt dictates the directional growth of CNS tissue along the central-peripheral axis

Finally, we investigated the effect of the Wnt/Tre1 signal-dependent orientation of NB polarity on the directional growth of the CNS tissue. In WT embryos, NBs are located at the peripheral-most part of the CNS tissue and generate progeny cells to the interior (more central region) of the embryo by divisions perpendicular to the overlying epithelium. This directional NB division allows the CNS tissue to grow along the central-peripheral axis of the *Drosophila* embryo, forming the layers of NBs, GMCs, and neurons from the peripheral to central direction (Fig. 6c and Supplementary Fig. 4c). Depletion of either Wnt or Tre1 signal resulted in the intermingling of NBs and their descendant cells, compromising the directional growth of the CNS tissue (Fig. 6d and e, 13.5 and 20.7% of NBs were located more centrally than their descendants, respectively, n=37 and 77; WT: 1.0%, n=96). Moreover, the simultaneous depletion of Wnt signals and *tre1* showed greater defects (Fig. 6f and Supplementary Fig. 4d, 40.7% of NBs were more centrally located than their descendants, n=118). Thus, Wnt cooperates with Tre1 to dictate the directional growth of the CNS tissue along the central-peripheral axis of the *Drosophila* embryo by controlling the orientation of NB polarity.

## Discussion

Here, we demonstrate that Wnt proteins from epithelial cells regulate the orientation of cell polarity in the adjacent NBs through a novel non-canonical Wnt signaling pathway and directly influence the apical polarity complex through fz and arm (Fig. 5g-g’’). Although arm is not essential for the formation of the apical polarity complex, its absence results in the abnormal positioning of the crescent of the polarity complex along the NB cortex. In contrast, the presence of arm promotes the formation of the trimeric complex consisting of arm, Baz, and Insc, and stabilizes the polarity complex at the Wnt-signaling side, thereby solidifying the relative orientation of Par-dependent polarity and asymmetric divisions. This orientation eventually directs the growth of the CNS tissue inward in the *Drosophila* embryo. Thus, Wnt proteins act as an instructive cue along the central-peripheral axis during *Drosophila* embryonic development.

The Wnt/fz/arm and Tre1-mediated signals appear to act independently in controlling the relative NB orientation in *Drosophila* embryos. This finding is consistent with the fact that these two signals operate through different entry points; the Par complex and the Pins complex, respectively. Indeed, when Wnt2 was supplied from *wg*-expressing cells in the *Wnt2/5/6/D* mutant (Fig. 2e), the NBs orientated between the direction of the Wnt2 signal and the apico-basal orientation, along which Tre1 signal is supposed to travel. Under these conditions, NB orientation may be determined by the sum of the Wnt2 and Tre1 signal vectors. Why these two simultaneous signals are required in NBs is unclear, and the molecules that activate the Tre1 signaling pathway remain to be identified. There may be other cases in which either of these signaling pathways solely determines the orientation of cell polarity or the division axis in response to extrinsic signals. Thus, these two mechanisms may have evolved independently to regulate cell behavior.

We also found that this non-canonical Wnt pathway facilitates the formation of a fully active Par complex in a non-cell-autonomous manner by enhancing the interaction between Baz and Insc. Although Insc has been primarily regarded as a bridge between the Par and Pins complexes, it appears that this protein has other functions in asymmetric NB division, as its linker function does not explain the defective localization of the cell fate determinants in *insc* mutants^21,23,29^. Although the detailed mechanisms are unclear, our results shed new light on this non-linker function of Insc. Available evidence suggests that Insc functions in various stem cell populations in various organisms^6,39-41^. The present results advance our understanding of the functions of Insc in the development and maintenance of tissue homeostasis.

Canonical Wnt signaling regulates various characteristics of stem cells involved in cell division, such as self-renewal^31,32^. Wnt proteins also regulate tissue polarity through non-canonical pathways, such as the PCP pathway^37,42^ which acts mostly within the epithelial plane. This study is the first to demonstrate that Wnt proteins regulate Par-dependent polarity orientation through a novel pathway. We also show that Wnt orients Par-dependent cell polarity orthogonally to the epithelial plane, perhaps through its secretion from the basal side of the epithelium. Wnt signaling thus regulates cell polarity in three dimensions relative to the epithelial structure, laterally through the PCP and orthogonally through Par-dependent polarity. Given that the epithelium is a fundamental structure in the formation and segregation of cell collectives, orientation of cell polarity by Wnt in multiple dimensions may play important roles in morphogenesis and/or tissue maintenance across species.

## Acknowledgements

We thank S. Hayashi, T. Kondo, T. Nishimura, N. Okamoto, T. Uemura, T. Usui, G. Struhl, M. Sato, C. Hama, T. Kojima, KYOTO Stock Center, and the Bloomington Stock Center for providing materials. We also thank T. Shibata for useful discussion.

## Contributions

S.Y. and F.M. designed the experiments, analyzed the data and wrote the manuscript.

S.Y. performed the experiments.

## Author information

The authors declare no competing financial interests.

## Methods

### Fly stocks

The *Wnt6* frame-shift mutants were constructed by the TALEN method^44,45^. TALE nucleases were designed to target exon 2 of the *Wnt6* gene as follows: Forward TCCAAATCTAATGTGCAAA and Reverse TTTCGGCCAACTTGCCGCG. Several lines of frame-shift mutants were recovered, including *Wnt6*^*Δ8-1*^ and *Wnt6*^*Δ28*^, which had 8-bp and 28-bp deletions, respectively: WT, TCCAAATCTAATGTGCAAAAAGACACGTCGTCTGCGCGGCAAGTTGGCCGAAA; *Wnt6*^*Δ8-1*^, TCCAAATCTAATGTGCAAAAAGA********CTGCGCGGCAAGTTGGCCGAAA;

*Wnt6*^*Δ28*^, T****************************GTCTGCGCGGCAAGTTGGCCGAAA. The *Df(2L)Wnt6-Wnt10* line was constructed by the recombination between *PBAC{WH}f02300* and *PBAC{WH}f05146*^46^.

*UAS-Wnt2-V5, UAS-Wnt5-V5, UAS-Wnt6-V5*, and *UAS-wntD-V5* were constructed by cloning the V5-tagged *Wnt* genes into *pUAST* vectors^47^.

Dominant negative forms of *fz* (*UAS-fz*^*ECD*^*-GFP*) were constructed by fusing a 1–273 amino acid fragment of *fz* with *GFP. UAS-fz2*^*ECD*^*-GPI* was used as a dominant negative *fz2*.

*UAS-arm* and *UAS-arm*^*XM19*^ were constructed by cloning WT and a 1–681 amino acid fragment of *arm* into *pUAST* vectors.

*w*^*1118*^ was used as WT.

The other fly lines were obtained from stock centers other than *Wnt5*^*D7*^ (C. Hama),

*fz3*^*G10*^ (T. Kojima), *fmi*^*E59*^, *fmi*^*E45*^ (T. Uemura) and *tre1*^*Δ10*13^. The following fly lines were used:

*Wnt2*^*O*^,

*Wnt5*^*D7*^,

*Wnt6*^*Δ8-1*^

*wntD*^*KO1*^

*Wnt5*^*D7*^; *Wnt2*^*O*^

*Wnt6*^*Δ8-1*^ *Wnt2*^*O*^

*Wnt2*^*O*^; *wntD*^*KO1*^

*Wnt5*^*D7*^; *Wnt6*^*Δ8-1*^

*Wnt5*^*D7*^; *wntD*^*KO1*^

*Wnt6*^*Δ8-1*^; *wntD*^*KO1*^

*Wnt5*^*D7*^; *Wnt6*^*Δ8-1*^ *Wnt2*^*O*^

*Wnt5*^*D7*^; *Wnt2*^*O*^; *wntD*^*KO1*^

*Wnt6*^*Δ28*^ *Wnt2*^*O*^; *wntD*^*KO1*^

*Wnt5*^*D7*^; *Wnt6*^*Δ8-1*^; *wntD*^*KO1*^

*Wnt5*^*D7*^; *Wnt6*^*Δ8-1*^ *Wnt2*^*O*^; *wntD*^*KO1*^

*Wnt5*^*D7*^; *Df(2L)BSC226 Wnt2*^*O*^; *wntD*^*KO1*^

*Wnt5*^*D7*^; *Insc-Gal4 Df(2L)Wnt6-Wnt10 Wnt2*^*O*^*/Wnt6*^*Δ8-1*^ *Wnt2*^*O*^; *UAS-Wnt2-V5 wntD*^*KO1*^*/wntD*^*KO1*^

*Wnt5*^*D7*^; *Insc-Gal4 Df(2L)Wnt6-Wnt10 Wnt2*^*O*^*/Wnt6*^*Δ8-1*^ *Wnt2*^*O*^; *UAS-Wnt5-V5 wntD*^*KO1*^*/wntD*^*KO1*^

*Wnt5*^*D7*^; *Insc-Gal4 Df(2L)Wnt6-Wnt10 Wnt2*^*O*^*/Wnt6*^*Δ8-1*^ *Wnt2*^*O*^; *UAS-Wnt6-V5 wntD*^*KO1*^*/wntD*^*KO1*^

*Wnt5*^*D7*^; *Insc-Gal4 Df(2L)Wnt6-Wnt10 Wnt2*^*O*^*/Wnt6*^*Δ8-1*^ *Wnt2*^*O*^; *UAS-wntD-V5 wntD*^*KO1*^*/wntD*^*KO1*^

*Wnt5*^*D7*^*/Y; Wnt6*^*Δ8-1*^ *Wnt2*^*O*^*/Wnt6*^*Δ28*^ *Wnt2*^*O*^; *UAS-Wnt2-V5 wntD*^*KO1*^*/wg-Gal4 wntD*^*KO1*^

*Wnt5*^*D7*^*/Y; Wnt6*^*Δ8-1*^ *Wnt2*^*O*^*/Wnt6*^*Δ28*^ *Wnt2*^*O*^; *UAS-wntD-V5 wntD*^*KO1*^*/wg-Gal4 wntD*^*KO1*^

*fz3*^*G10*^ *fz4*^*3-1*^*/Y; Insc-Gal4/UAS-fz*^*ECD*^*-GFP UAS-fz2*^*ECD*^*-GPI*

*fz3*^*G10*^*/Y; Insc-Gal4/UAS-fz*^*ECD*^*-GFP UAS-fz2*^*ECD*^*-GPI*

*fz4*^*3-1*^*/Y; Insc-Gal4/UAS-fz*^*ECD*^*-GFP UAS-fz2*^*ECD*^*-GPI*

*fz3*^*G10*^ *fz4*^*3-1*^*/Y; Insc-Gal4/UAS-fz*^*ECD*^*-GFP*

*fz3*^*G10*^ *fz4*^*3-1*^*/Y; Insc-Gal4/UAS-fz2*^*ECD*^*-GPI*

*arm*^*1*^

*da-Gal4/UAS-Axin-GFP*

*arm*^*1*^*/Y; Insc-Gal4/+; UAS-arm/+*

*pan*^*2*^

*arm*^*1*^*/Y; Insc-Gal4/+; UAS-arm*^*XM19*^*/+*

*fmi*^*E59*^

*fmi*^*E45*^

*Vang*^*stbm-6*^

*dgo*^*380*^

*pk*^*sple-1*^

*arm*^*2*^ *FRT101 tre1*^*Δ10*^

*arm*^*2*^ *tre1*^*Δ10*^ *FRT9-2*

*fz3*^*G10*^ *tre1*^*Δ10*^ *fz4*^*3-1*^*/Y; Insc-Gal4/UAS-fz*^*ECD*^*-GFP UAS-fz2*^*ECD*^*-GFP*

*tre1*^*Δ10*^*/Y; arm-Gal4/+; UAS-Axin-GFP/+*

*Insc-Gal4/+; UAS-Axin-GFP/+*

*tre1*^*Δ10*^*/Y; Insc-Gal4/+; UAS-Axin-GFP/+*

### Immunohistochemistry

Immunohistochemical staining of *Drosophila* embryos was performed as described previously^13^.

For arm staining, antigen retrieval was performed using HistoVT One (nacalai tesque). For preparing *Drosophila* embryo sections, stained embryos were embedded in OCT compound (Sakura Finetek) and cut on a cryostat.

The following antibodies were used: mouse anti-Mira^48^, rabbit anti-Mira^16^, rabbit anti-pH3 (Upstate, 06-570), mouse anti-En (DSHB, 4D9), mouse anti-wg (DSHB, 4D4), mouse anti-Wor (X. Yang, IMCB), rat anti-Pros^49^, rabbit anti-Baz^48^, rabbit anti-aPKC (Santa Cruz, C-20), rabbit anti-Par-6^50^, rabbit anti-Pins^51^, rabbit anti-Insc^13^, guinea pig anti-Centrosomin^13^, mouse anti-arm (DSHB, N2 7A1), chicken anti-GFP (Aves labs, GFP-1020), chicken anti-beta-Galactosidase (abcam, ab9361), and mouse anti-Sxl (DSHB, M18).

### Fluorescent *in situ* hybridization

Fluorescent *in situ* hybridization was performed as previously described^52^.

Templates for the probes for *Wnt* genes were amplified by PCR using the following primers: *wg*-F;ATGGATATCAGCTATATCTTCGTCATCTGC, *wg*-R; AGACACGTGTAGATGACCTTTTTGGTCC, *Wnt2*-F; ATGTGGAAAATACATAACAAGC, *Wnt2*-R; CCTTTACAGGTGAACTCCTCGTAGCTCTCG, *Wnt5*-F; ATGAGTTGCTACAGAAAAAGG, *Wnt5*-R; CCTTTACATGTGTGCTCCTCGAGTACC, *Wnt6*-F; ATGCGTTTGCTCATGGTAATTGC, *Wnt6*-R; CCGAGGCAGGTGTTGACCGCCCGGTGTTCC, *Wnt10*-F; ATGACTGCGTGGCGAGCAACATCAAAAGG, *Wnt10*-R; CCATTGCATATGCTAATCCATTCCTCTACG, *wntD*-F; ATGATTTTTGCCATCACATTCTTCATGG and *wntD*-R; GTAGCAGGAGTACTGCCTTTCCAGCTGG.

### Measurement of the NB division angle in 3D

Measurement of the NB orientation was performed as previously described^43^.

### Co-immunoprecipitation of *Drosophila* S2 cell lysate

The maintenance and transfection of S2 cells were performed as previously described^13^. Cells were suspended in lysis buffer [20 mM Tris-HCl (pH7.5), 150 mM NaCl, 1 mM EDTA, 10% glycerol, 0.5% NP-40, Protease Inhibitor Cocktail and Phosphatase Inhibitor Cocktail (nacalai tesque)]. The cleared cell lysate was subjected to immunoprecipitation using a rabbit anti-Flag antibody (SIGMA, F7425) or mouse anti-Myc antibody (Millipore, 05-724) with Dynabeads Protein G (VERITAS), and the immunoprecipitate was probed with rabbit anti-Flag (SIGMA, F7425), mouse anti-arm (DSHB, N2 7A1), or chicken anti-Myc antibodies (BETHYL, A190-103A).

### Co-immunoprecipitation of *Drosophila* embryo lysates

Co-immunoprecipitation of *Drosophila* embryonic lysates was performed as previously described^13^.

Embryos were suspended in lysis buffer as described above. The cleared cell lysate was subjected to immunoprecipitation using a rabbit anti-Baz antibody^48^, rabbit anti-Insc antibody^13^, or normal rabbit IgG (MBL, PM035) with Dynabeads Protein G (VERITAS), and the immunoprecipitate was probed with rabbit anti-Baz^48^, rabbit anti-Insc^13^, or mouse anti-arm antibodies (DSHB, N2 7A1).

### Statistics

Data were analyzed in Prism 6 (GraphPad). Data were compared by two-tailed, unpaired Student’s *t*-tests unless otherwise noted. No statistical methods were used to predetermine sample size. The experiments were not randomized, and the investigators were not blinded to allocation during experiments and outcome assessment.

## References

1. Baena-López, L. A., Baonza, A. & García-Bellido, A. The orientation of cell divisions determines the shape of Drosophila organs. Current Biology 15, 1640–1644 (2005).

2. Fischer, E. et al. Defective planar cell polarity in polycystic kidney disease. Nat Genet 38, 21–23 (2005).

3. Tang, N., Marshall, W. F., McMahon, M., Metzger, R. J. & Martin, G. R. Control of mitotic spindle angle by the RAS-regulated ERK1/2 pathway determines lung tube shape. Science 333, 342–345 (2011).

4. Yamashita, Y. M., Jones, D. L. & Fuller, M. T. Orientation of asymmetric stem cell division by the APC tumor suppressor and centrosome. Science 301, 1547–1550 (2003).

5. Xiong, F. et al. Interplay of Cell Shape and Division Orientation Promotes Robust Morphogenesis of Developing Epithelia. Cell 159, 415–427 (2014).

6. Williams, S. E., Ratliff, L. A., Postiglione, M. P., Knoblich, J. A. & Fuchs, E. Par3–mInsc and Gαi3 cooperate to promote oriented epidermal cell divisions through LGN. Nature Cell Biology 16, 758–769 (2014).

7. Kamakura, S. et al. The Cell Polarity Protein mInscRegulates Neutrophil Chemotaxisvia a Noncanonical G Protein Signaling Pathway. Developmental Cell 26, 292–302 (2013).

8. Goulas, S., Conder, R. & Knoblich, J. A. The Par complex and integrins direct asymmetric cell division in adult intestinal stem cells. Cell Stem Cell 11, 529–540 (2012).

9. Arata, Y., Lee, J.-Y., Goldstein, B. & Sawa, H. Extracellular control of PAR protein localization during asymmetric cell division in the C. elegans embryo. Development 137, 3337–3345 (2010).

10. Knoblich, J. A. Mechanisms of Asymmetric Stem Cell Division. Cell 132, 583–597 (2008).

11. Prehoda, K. E. Polarization of Drosophila Neuroblasts During Asymmetric Division. Cold Spring Harbor Perspectives in Biology 1, a001388–a001388 (2009).

12. Yu, F., Kuo, C. T. & Jan, Y.-N. Drosophila neuroblast asymmetric cell division: recent advances and implications for stem cell biology. Neuron 51, 13–20 (2006).

13. Yoshiura, S., Ohta, N. & Matsuzaki, F. Tre1 GPCR Signaling Orients Stem Cell Divisions in the Drosophila Central Nervous System. Developmental Cell 22, 79–91 (2012).

14. Betschinger, J., Mechtler, K. & Knoblich, J. A. Asymmetric Segregation of the Tumor Suppressor Brat Regulates Self-Renewal in Drosophila Neural Stem Cells. Cell 124, 1241–1253 (2006).

15. Hirata, J., Nakagoshi, H., Nabeshima, Y. & Matsuzaki, F. Asymmetric segregation of the homeodomain protein Prospero during Drosophila development. Nature 377, 627–630 (1995).

16. Ikeshima-Kataoka, H., Skeath, J. B., Nabeshima, Y., Doe, C. Q. & Matsuzaki, F. Miranda directs Prospero to a daughter cell during Drosophila asymmetric divisions. Nature 390, 625–629 (1997).

17. Knoblich, J. A., Jan, L. Y. & Jan, Y. N. Asymmetric segregation of Numb and Prospero during cell division. Nature 377, 624–627 (1995).

18. Lu, B., Rothenberg, M., Jan, L. Y. & Jan, Y. N. Partner of Numb colocalizes with Numb during mitosis and directs Numb asymmetric localization in Drosophila neural and muscle progenitors. Cell 95, 225–235 (1998).

19. Petronczki, M. & Knoblich, J. A. DmPAR-6 directs epithelial polarity and asymmetric cell division of neuroblasts in Drosophila. Nature Cell Biology 3, 43–49 (2001).

20. Rhyu, M. S., Jan, L. Y. & Jan, Y. N. Asymmetric distribution of numb protein during division of the sensory organ precursor cell confers distinct fates to daughter cells. Cell 76, 477–491 (1994).

21. Schober, M., Schaefer, M. & Knoblich, J. A. Bazooka recruits Inscuteable to orient asymmetric cell divisions in Drosophila neuroblasts. Nature 402, 548–551 (1999).

22. Shen, C. P., Jan, L. Y. & Jan, Y. N. Miranda is required for the asymmetric localization of Prospero during mitosis in Drosophila. Cell 90, 449–458 (1997).

23. Wodarz, A., Ramrath, A., Kuchinke, U. & Knust, E. Bazooka provides an apical cue for Inscuteable localization in Drosophila neuroblasts. Nature 402, 544–547 (1999).

24. Wodarz, A., Ramrath, A., Grimm, A. & Knust, E. Drosophila atypical protein kinase C associates with Bazooka and controls polarity of epithelia and neuroblasts. The Journal of Cell Biology 150, 1361–1374 (2000).

25. Parmentier, M. L. et al. Rapsynoid/partner of inscuteable controls asymmetric division of larval neuroblasts in Drosophila. J. Neurosci. 20, RC84 (2000).

26. Schaefer, M., Shevchenko, A., Shevchenko, A. & Knoblich, J. A. A protein complex containing Inscuteable and the Galpha-binding protein Pins orients asymmetric cell divisions in Drosophila. Curr. Biol. 10, 353–362 (2000).

27. Schaefer, M., Petronczki, M., Dorner, D., Forte, M. & Knoblich, J. A. Heterotrimeric G proteins direct two modes of asymmetric cell division in the Drosophila nervous system. Cell 107, 183–194 (2001).

28. Yu, F., Morin, X., Cai, Y., Yang, X. & Chia, W. Analysis of partner of inscuteable, a novel player of Drosophila asymmetric divisions, reveals two distinct steps in inscuteable apical localization. Cell 100, 399–409 (2000).

29. Kraut, R., Chia, W., Jan, L. Y., Jan, Y. N. & Knoblich, J. A. Role of inscuteable in orienting asymmetric cell divisions in Drosophila. Nature 383, 50–55 (1996).

30. Siegrist, S. E. & Doe, C. Q. Extrinsic cues orient the cell division axis in Drosophila embryonic neuroblasts. Development 133, 529–536 (2006).

31. Nusse, R. et al. Wnt signaling and stem cell control. Cold Spring Harb. Symp. Quant. Biol. 73, 59–66 (2008).

32. Cadigan, K. M. & Peifer, M. Wnt Signaling from Development to Disease: Insights from Model Systems. Cold Spring Harbor Perspectives in Biology 1, a002881–a002881 (2009).

33. Huang, H.-C. & Klein, P. S. The Frizzled family: receptors for multiple signal transduction pathways. Genome Biology 5, 234 (2004).

34. Miller, J. R. The Wnts. Genome Biology 3, REVIEWS3001 (2002).

35. Willert, K., Logan, C. Y., Arora, A., Fish, M. & Nusse, R. A Drosophila Axin homolog, Daxin, inhibits Wnt signaling. Development 126, 4165–4173 (1999).

36. Chan, S.-K. & Struhl, G. Evidence that Armadillo transduces wingless by mediating nuclear export or cytosolic activation of Pangolin. Cell 111, 265–280 (2002).

37. Devenport, D. The cell biology of planar cell polarity. The Journal of Cell Biology 207, 171–179 (2014).

38. Wei, S.-Y. et al. Echinoid Is a Component of Adherens Junctions That Cooperates with DE-Cadherin to Mediate Cell Adhesion. Developmental Cell 8, 493–504 (2005).

39. Žigman, M. et al. Mammalian Inscuteable Regulates Spindle Orientation and Cell Fate in the Developing Retina. Neuron 48, 539–545 (2005).

40. Postiglione, M. P. et al. Mouse Inscuteable Induces Apical-Basal Spindle Orientation to Facilitate IntermediateProgenitor Generation in the Developing Neocortex. Neuron 72, 269–284 (2011).

41. Ballard, M. S. et al. Mammary Stem Cell Self-Renewal Is Regulated by Slit2/Robo1 Signaling through SNAI1 and mINSC. CellReports (2015). doi:10.1016/j.celrep.2015.09.006

42. Sugioka, K., Mizumoto, K. & Sawa, H. Wnt Regulates Spindle Asymmetry to Generate AsymmetricNuclear b-Catenin in C. elegans. Cell 146, 942–954 (2011).

43. Jüschke, C., Xie, Y., Postiglione, M. P. & Knoblich, J. A. Analysis and modeling of mitotic spindle orientations in three dimensions. Proc. Natl. Acad. Sci. U.S.A. 111, 1014–1019 (2014).

44. Sakuma, T. et al. Efficient TALEN construction and evaluation methods for human cell and animal applications. Genes to Cells 18, 315–326 (2013).

45. Kondo, T. et al. TALEN-induced gene knock out in Drosophila. Dev Growth Differ 56, 86–91 (2014).

46. Parks, A. L. et al. Systematic generation of high-resolution deletion coverage of the Drosophila melanogaster genome. Nat Genet 36, 288–292 (2004).

47. Brand, A. H. & Perrimon, N. Targeted gene expression as a means of altering cell fates and generating dominant phenotypes. Development 118, 401–415 (1993).

48. Ohshiro, T., Yagami, T., Zhang, C. & Matsuzaki, F. Role of cortical tumour-suppressor proteins in asymmetric division of Drosophila neuroblast. Nature 408, 593–596 (2000).

49. Fuse, N., Hisata, K., Katzen, A. L. & Matsuzaki, F. Heterotrimeric G proteins regulate daughter cell size asymmetry in Drosophila neuroblast divisions. Current Biology 13, 947–954 (2003).

50. Izumi, Y., Ohta, N., Itoh-Furuya, A., Fuse, N. & Matsuzaki, F. Differential functions of G protein and Baz-aPKC signaling pathways in Drosophila neuroblast asymmetric division. The Journal of Cell Biology 164, 729–738 (2004).

51. Izumi, Y., Ohta, N., Hisata, K., Raabe, T. & Matsuzaki, F. Drosophila Pins-binding protein Mud regulates spindle-polarity coupling and centrosome organization. Nature Cell Biology 8, 586–593 (2006).

52. Nagaso, H., Murata, T., Day, N. & Yokoyama, K. K. Simultaneous detection of RNA and protein by in situ hybridization and immunological staining. J. Histochem. Cytochem. 49, 1177–1182 (2001).

