## Supplemental figures for "A novel Wnt pathway orients Par complex-dependent cell polarity"

a

| Mutant/Deficiency lines | Deleted genes | UAS lines | Expressed gene |
|---|---|---|---|
| *Df(1)ED6712* | *Ilp7* | *P{XP}dlpd10506* | *dlp* |
| *Df(2L)BSC226* | *wg, Wnt4, Wnt6, Wnt10* | *P{XP}d03132* | *Wnt2* |
| *Df(2L)Exel6017* | *wg, Wnt4, Wnt6, Wnt10* | *P{GSV6}GS11907* | *Ten-m* |
| *Df(2L)BSC299* | *CG31832* | *P{GSV1}GS1209* | *trol* |
| *Df(2R)Exel7170* | *CG30280* | *P{GSV1}Wnt5GS1192* | *Wnt5* |

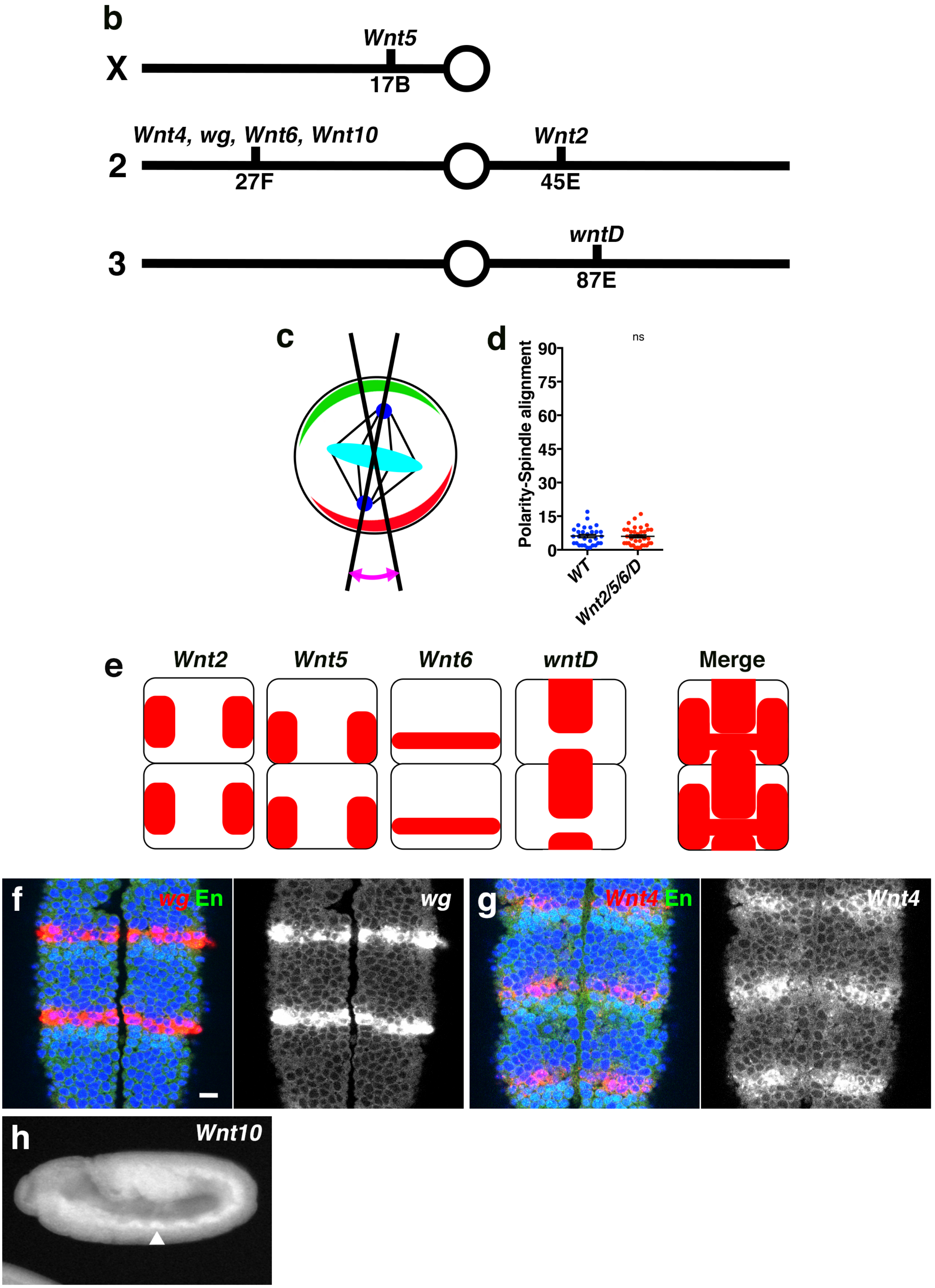


sFig. 1

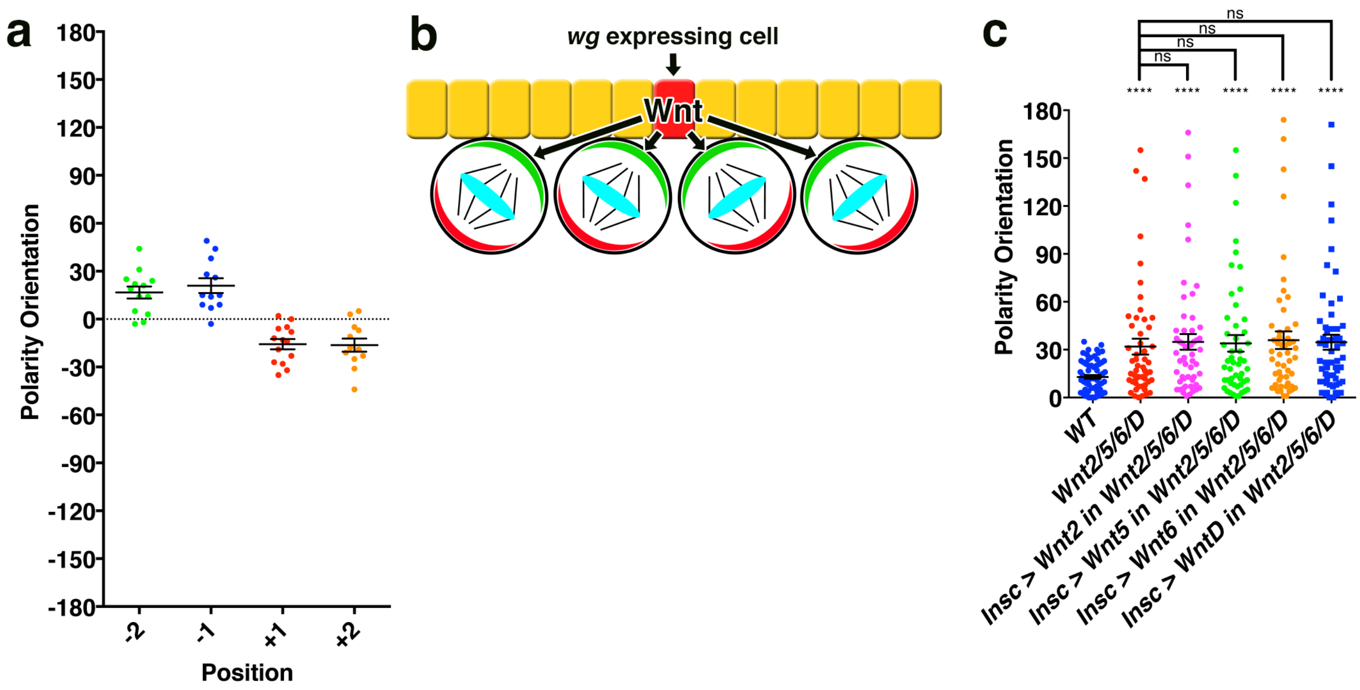
a
Polarity Orientation
180
150
120
90
60
30
0
-30
-60
-90
-120
-150
-180
-2
-1
+1
+2
Position
b
wg expressing cell
Wnt
c
ns
ns
ns
ns
****
****
****
****
****
Polarity Orientation
180
150
120
90
60
30
0
WT
Wnt2/5/6/D
Insc > Wnt2 in Wnt2/5/6/D
Insc > Wnt5 in Wnt2/5/6/D
Insc > Wnt6 in Wnt2/5/6/D
Insc > WntD in Wnt2/5/6/D

sFig. 2

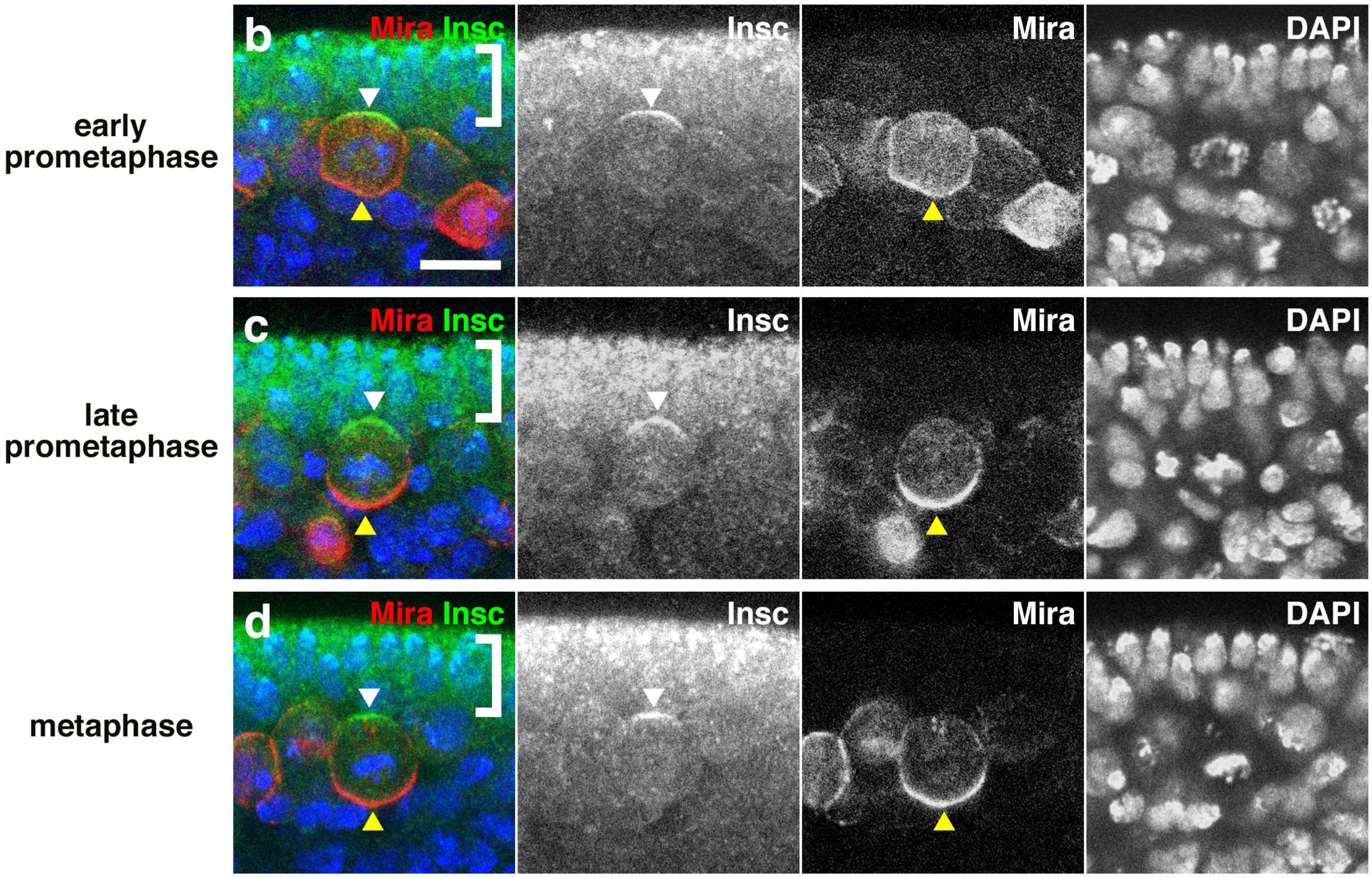
early prometaphase
late prometaphase
metaphase
b
c
d
Mira Insc
Insc
Mira
DAPI

sFig. 3

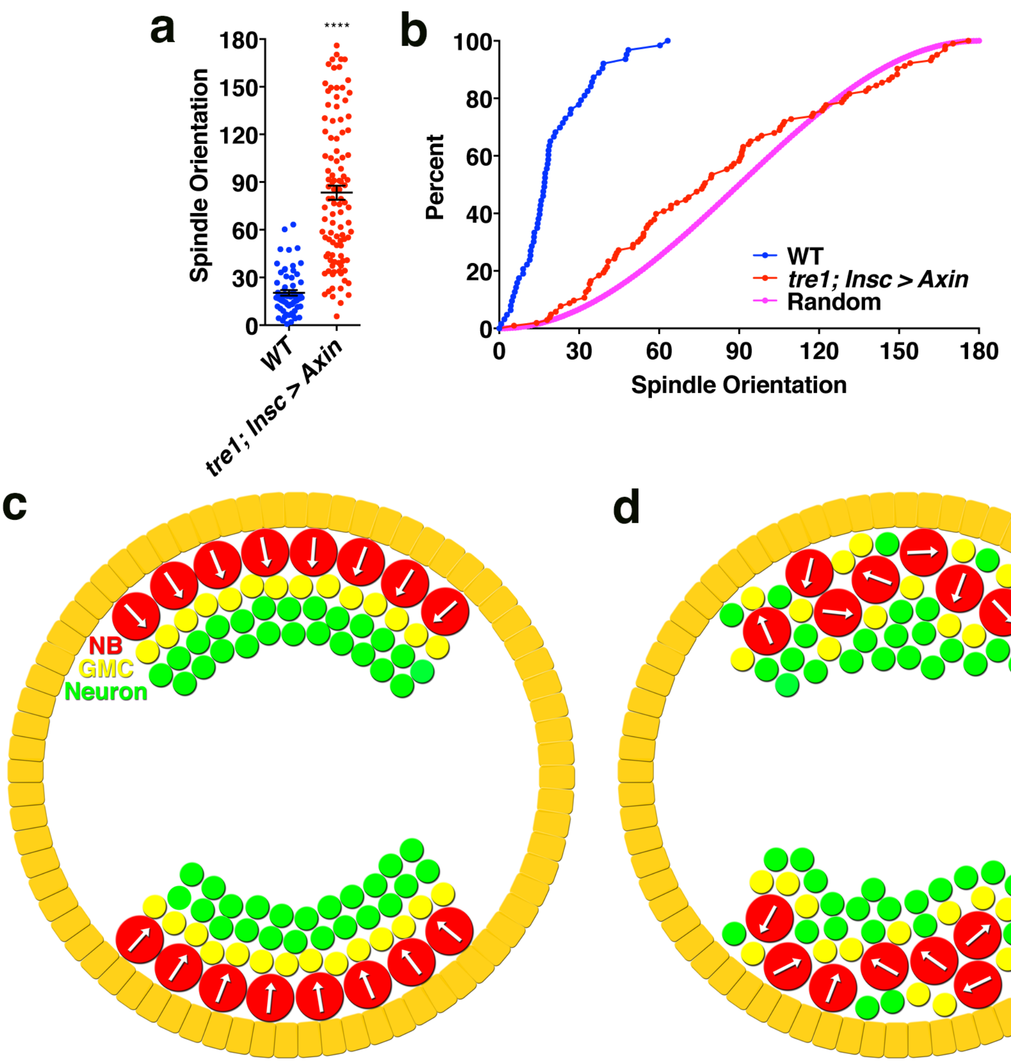
a
Spindle Orientation
180
150
120
90
60
30
0
****
WT
tre1; Insc > Axin
b
Percent
100
80
60
40
20
0
0
30
60
90
120
150
180
Spindle Orientation
WT
tre1; Insc > Axin
Random
c
NB
GMC
Neuron
d

sFig.. 4

### Supplementary Figure legends

**Supplementary Figure 1. Wnt2, Wnt5, Wnt6, and wntD redundantly regulate the orientation of NB polarity.**

(a) Candidate lines for the signaling molecules obtained by the LOF and GOF screens. Mutant/deficiency lines used in the LOF screen and UAS lines used in the GOF screen are shown. Primary candidate genes in the LOF screen are shown as "deleted genes". (b) Schematic drawing of the cytological locations of *Wnt* genes in the *Drosophila* genome. (c) Schematic drawing of the method used to measure the alignment between NB orientation and spindle axis. Red and green crescents indicate Mira and Par complex localization, respectively, and a blue circle indicates the position of the centrosome as visualized using centrosomin staining. Angle between the Par complex polarity orientation and spindle axis was measured. (d) The alignment between the Par complex orientation and spindle axis in WT and *Wnt2/5/6/D* mutant (angle). Mean±SEM values are shown in the plots. Sample number *n*=30, 36 (from left to right). The P value is 0.89. (e) Schematic drawing of the expression patterns of *Wnt2*, *Wnt5*, *Wnt6*, and *wntD*. Dorsal views of the epithelial layers overlying the CNS region are shown. (f-h) Expression patterns of *wg* (f), *Wnt4* (g), and *Wnt10* (h). *Wnt10* is expressed in the mesoderm (arrowhead). (f and g): Dorsal view, anterior is up. (h): Lateral view, anterior is left. Scale bar: 10 μm.

**Supplementary Figure 2. Wnt proteins from epithelial cells act as directional cues to orient NB polarity.**

(a) NB orientation in *Wnt2/5/6/D* mutants expressing *wntD* from *wg-Gal4*. The angle of the NB orientation relative to the epithelial plane was defined as described in the Fig. 1 legend. The mean angles are as follows: position–2: 16.7°, position–1: 20.9°, position+1: –15.7°, position+2: –16.3°. Mean±SEM values are shown in the plots. The individual sample number *n*=13, 12, 14, 12 (from left to right). (b) Schematic drawing of the NB orientation in embryos expressing *WntD* from *wg-Gal4* in a *Wnt2/5/6/D* mutant background (see text for details). A red cell indicates a wg-positive cell that expresses *WntD* (c) The NB orientation of *Wnt2/5/6/D* mutant embryos expressing one of the 4 effective Wnts in NBs and their progenies. No rescue of NB orientation was observed. Mean±SEM is shown in the plots. The data for WT, *Wnt2/5/6/D*, and *Insc* >

*Wnt6* in the *Wnt2/5/6/D* mutant background are the same as those shown in Fig. 2f. ns: not significant. ****: $p<0.0001$. The individual sample number n and P value (compared with *Wnt2/5/6/D*) in the parenthesis are as follows (from left to right): $n=70$, 52, 55 (0.67), 50 (0.78), 52 (0.58), 58 (0.69).

**Supplementary Figure 3. Time course of the basal localization of Mira during the mitotic phase of NBs.**

Basal localization of Mira started at the end of interphase and was not completed during early prometaphase (a). Mira localization was completed at late prometaphase (b) and persisted through metaphase (c). Apical localization of polarity proteins such as Insc was completed at the end of interphase and persisted through prometaphase and metaphase (a-c). Localization of Mira and Insc are indicated by yellow and white arrowheads, respectively. Brackets: epithelial layer. Scale bar: 10 μm.

**Supplementary Figure 4. Wnt/fz/arm signal regulates NB orientation and directional CNS tissue growth in parallel with Tre1.**

(a and b) Statistics of the NB division angles in WT embryos, and *tre1*mutant embryos expressing Axin in NBs and their progenies. Spindle orientation was measured in the 3-D way[43]. Centrosomin was used as a marker for the spindle orientation. The Mean±SEM values are shown in in the plots (a). The sample number $n=63$ for WT, 103 for *Tre1; Insc>Axin*. The P value is $< 0.0001$. Dot plots (a) and cumulative graphs (b) were prepared from the same data sets. (c and d) Schematic drawing of the directional growth of the CNS tissue along the central-peripheral axis of the *Drosophila* embryo in WT (c) and a *tre1* mutant expressing *Axin* in the NBs (d). Red: positions of NB. Yellow: positions of GMCs. Green: positions of neurons. Orange: epithelial cells.
